# Influence of anthropogenic environments on the activity patterns and vigilance of Cape chacma baboons (*Papio ursinus ursinus*) in the Garden Route, South Africa

**DOI:** 10.1101/2025.11.17.688758

**Authors:** S. Dany, C. Lacomme, A. Guionneau, A. Ramaru, F. Prugnolle, O. Petit, V. Rougeron

## Abstract

Anthropogenic activities shape animals time allocation and risk perception, yet its immediate behavioural impacts on primate species remain poorly quantified. In the present study, variation in activity budgets and vigilance were analysed in three Cape chacma baboon (*Papio ursinus ursinus*) troops inhabiting contrasting anthropogenic environments in South Africa’s Garden Route: Madiba (in a peri-urban campus), North (in an intermediate area), and Outeniqua (in a more rural area). From March to June 2024, a total 84 h of observations were recorded yielding 343 scan samples, and modelled behaviour as a function of troop and environmental covariates (weather, habitat openness, car and human presence). Feeding dominated overall time allocation (35.4%), with no whole-budget differences among troops, but behaviour-specific contrasts emerged. Outeniqua troop inhabiting the more rural area relied significantly less on human-derived foods than Madiba and showed less running than both Madiba and North. In addition, level-3 vigilance (active/intense scanning) was higher in the troops frequenting more anthropogenic environments (Madiba, North) than in Outeniqua, suggesting that vigilance is a particularly sensitive response to human exposure. Finally, habitat openness was negatively associated with affiliative interactions, and car presence was negatively associated with resting. These results provide partial support for hypothesised disturbance effects, with shifts in behavioural sub-categories without overall budget change, clarifying how anthropogenic disturbance can restructure primate time allocation and risk perception. Such immediate behavioural responses should offer actionable insight for evidence-based conservation and human–baboon conflict mitigation.

## Introduction

Human activities expansion stands as one of the leading driver of wildlife extinction, mainly through the fragmentation, degradation, and loss of natural habitats and resources (Vitousek et al. 1997; Guy Cowlishaw & Robin I. M. Dunbar 2000; Coppolillo 2005; Estrada et al. 2017; Tilman et al. 2017). The worldwide constant growth of anthropogenic landscapes forces an increasing number of species to adapt to human-modified environments (Achard et al. 2001; Muche et al. 2023). This transition introduces new ecological and behavioural challenges, requiring species to adapt to survive in habitats increasingly dominated by human presence (Hulme-Beaman et al. 2016). Generalist diets, high behavioural flexibility, and tolerance to human disturbance tend to promote persistence, whereas strong habitat specialization and low diet plasticity increase vulnerability, contributing to pronounced population declines under fragmentation and disturbance (Prange, S. and Gehrt, S. D. and Wiggers 2004 Jun; LaDeau et al. 2013; Maisels et al. 2013; Tempel et al. 2014; Dugger et al. 2016; Murray and St. Clair 2017). Species with slower life histories (i.e. longer lifespans, delayed reproduction, fewer offspring, and greater parental investment) are less resilient to rapid environmental change because population recovery is slow and disturbance effects accumulate across generations. In addition to this demographic sensitivity, some taxa face additive direct pressures where their space-use or foraging brings them into frequent conflict with humans (e.g. crop-raiding, infrastructure damage, perceived risk to safety) (Larson et al. 2016; Schell et al. 2021). Such conflict elevates mortality and displacement through deterrence, translocation, or lethal control, making these species particularly vulnerable in human-dominated landscapes, even when their life-history pace alone would not predict extreme sensitivity (as in many large herbivores and behaviourally bold primates) (i.e. Larson et al. 2016). By contrast, species with faster life histories and high behavioural flexibility can exploit novel resources and microhabitats created by people, allowing persistence, or even proliferation, under intensive landscape modification (Hetmański et al. 2011; Bateman and Fleming 2012; Scholz et al. 2020). Such species exhibit broad ecological tolerance and high behavioural plasticity, allowing rapid adjustment to novel conditions (Charmantier and Gienapp 2014; Sih et al. 2016). As global environmental changes accelerate, understanding interspecific differences in adaptive capacity has become crucial for preserving biodiversity in increasingly human-dominated ecosystems.

Among mammals, primates represent a particularly informative model for studying behavioural responses to anthropogenic environments, due to their cognitive complexity, ecological flexibility, and evolutionary proximity to humans (McLennan et al. 2017; McLennan et al. 2020; Bersacola et al. 2023). While anthropogenic habitats can offer certain benefits, such as abundant, high-energy food resources and reduced predation risk, they also impose costs that alter natural behaviours, including shifts in diet, movement, and exploratory activity (Saj et al. 2001; Kaplan et al. 2011; Fehlmann et al. 2017; Corrêa et al. 2018; Katlam et al. 2018; Dhananjaya et al. 2022; Kennedy Overton et al. 2024). Examples illustrating this behavioural plasticity in primates are widespread. For instance, the snow monkeys (*Macaca fuscata*) in Japan and chacma baboons (*Papio ursinus ursinus*) in South Africa incorporate substantial amounts of human-provided food into their diets, such as processed carbohydrates and protein-rich items (Hoffman and O’Riain 2012; Oi et al. 2021; Kennedy Overton et al. 2024). In Bali, Indonesia, long-tailed macaques (*Macaca fascicularis*) have adapted to urban settings by actively exploring for food among tourists, including searching through bags and entering buildings (Riley and Fuentes 2011), while Samango monkeys (*Cercopithecus albogularis*) in South Africa display risk-sensitive foraging near dwellings (Novak et al. 2014). Other species, including olive baboons (*Papio anubis*) in Kenya, Barbary macaques (*Macaca sylvanus*) in Morocco, vervet monkeys (*Chlorocebus pygerythrus*) in Uganda, and bonnet macaques (*Macaca radiata*) in India, have all shown altered activity budgets when confronting to anthropogenic environments, spending more time near settlements and altering time allocations for grooming, foraging, and movement (Kumara et al. 2010; Sha and Hanya 2013; Novak et al. 2014; Lewis and O’Riain 2017; Thatcher et al. 2020). Collectively, these behavioural shifts highlight the ecological and evolutionary consequences of anthropogenic changes for primates and highlight the importance of understanding how human presence modifies their activity patterns and social dynamics, particularly in human-conflict areas (Almeida-Rocha et al. 2017).

Across sub-Saharan Africa, *Papio* is among the most successful and widely distributed primate genera, inhabiting a diverse range of habitats, climates, and latitudes (Henzi and Barrett 2003). This success is largely attributed to their remarkable adaptability, which includes a highly flexible omnivorous diet, complex social organisation, variable mating systems, and versatile locomotion (Alberts and Altmann; Whiten et al. 1987; Else 1991; Whiten, A. and Byrne, R. W. and Barton, R. A. and Waterman, P. G. and Henzi 1991; Henzi and Barrett 2003). Among its six species, the chacma baboon, *Papio ursinus*, occupies the African southernmost range in a mosaic of natural and anthropogenic landscapes (Whiten et al. 1987; Henzi and Barrett 2003). Its proximity to humans has, however, led to growing conflict, as baboons raid crops and human waste or enter homes for food (Hill 2000; Chowdhury et al. 2020; Kennedy Overton et al. 2024). Although chacma baboons are legally protected under the Western Cape Nature Conservation Act (2000), chacma baboons are often killed in retaliation, through poisoning or shooting (Beamish and O’Riain 2014). Prior studies have documented how chacma baboons adapt to anthropogenic environments by exploiting novel food resources and undergoing changes in their social relationships (Henzi et al. 2011; Pebsworth et al. 2012; Lewis and O’Riain 2017; Mazué et al. 2023). However, relatively little is known about how anthropogenic environments affect core behaviours such as vigilance and the allocation of time to key activities. These metrics provide valuable insight into behavioural adaptation and perceived risk, reflecting both immediate pressures and longer-term strategies. The current scarcity of comparative, multi-troop studies likely reflects the logistical difficulty of monitoring free-ranging baboons across heterogeneous landscapes.

The present study seeks to address this gap by analysing activity budgets and vigilance in three chacma baboon troops inhabiting contrasting anthropogenic environments in the Garden Route region of South Africa. Over a three months period (March to June 2024), we monitored: (i) the Madiba troop sharing a peri-urban campus, (ii) the North troop, inhabiting an environment that lies between rural and peri-urban campus settings, and (iii) the Outeniqua troop, in a predominantly rural environment (Fig. 1). We evaluated how these troops allocated time among key behaviours, including affiliation, agonistic, feeding, locomotion, resting, sexual and vigilance. We formulated three hypotheses: *(H1) Anthropogenic disturbance reshapes baboon activity budgets via two non-exclusive pathways*. Where access to predictable, high-energy human-derived foods and low deterrence, reduced foraging effort will free time that is reallocated to low-cost behaviours (e.g., resting, affiliation). Where human and car are present, saved foraging time will be offset by greater vigilance and locomotion behaviors; (H2) *North troop, inhabiting an intermediate setting, should exhibit activity patterns and vigilance levels intermediate between rural and peri-urban troops*, reflecting a gradient of behavioural adaptation to human presence; (H3) *Anthropogenic activity (human and car presence) should increase vigilance in the peri-urban Madiba troop, in comparison to the two other troops inhabiting less anthropic areas*. To complement these hypotheses, we tested environmental covariates, including weather, habitat openness, cars and human presence, in order to identify whether they mediate the behavioural patterns predicted above. This study aims to provide new insights into how chacma baboons adjust core behavioural strategies in response to human disturbance and environmental change, thereby contributing to broader understanding of primate resilience in an increasingly urbanized world.

**Figure 1.**
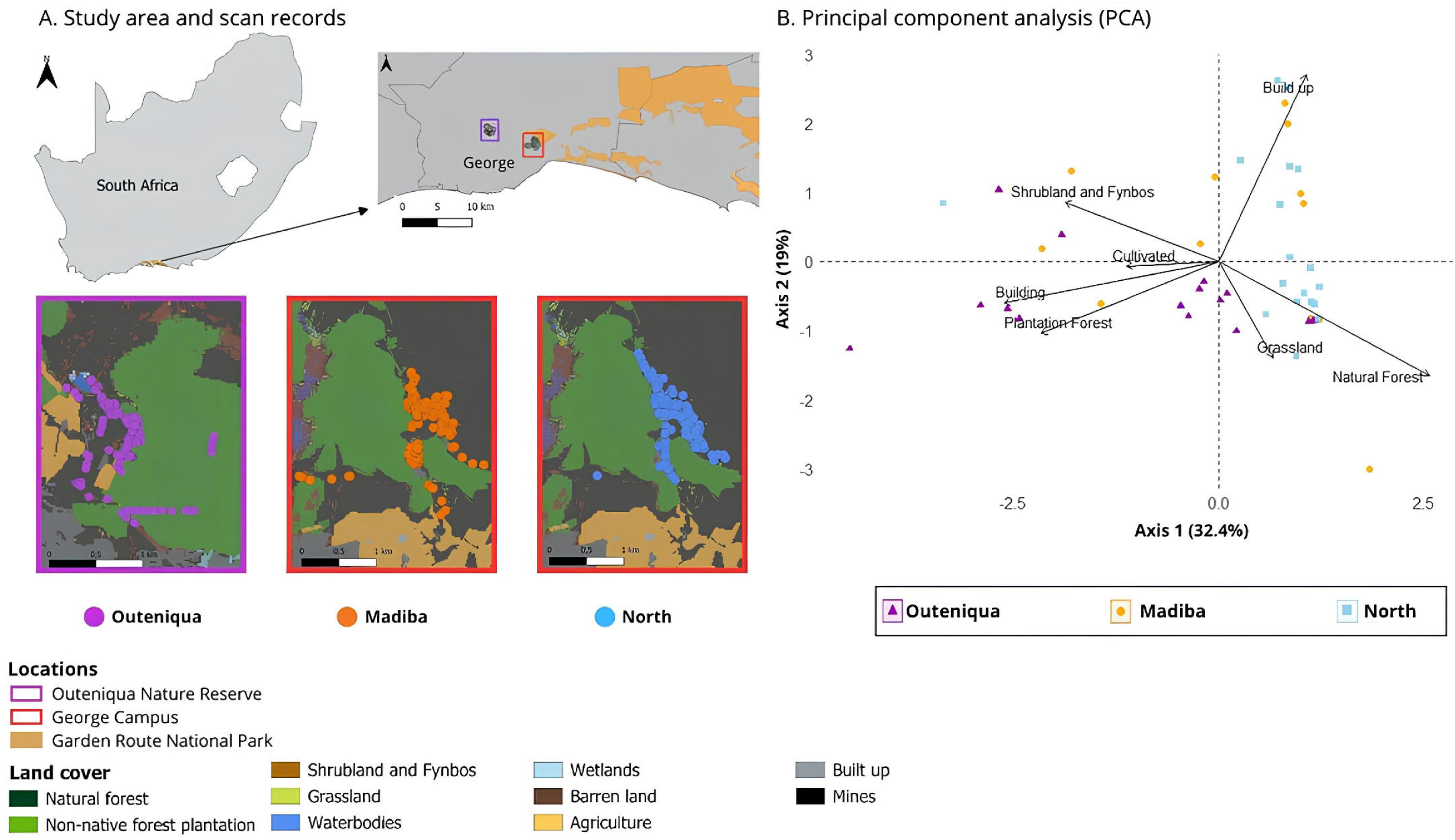
Spatial distribution of study sites and land use context of chacma baboon scan data in the Garden Route, South Africa. A. Study area and scan records in the Garden Route, Western Cape, South Africa. The top left map displays the study site within the Garden Route region of South Africa. The top right map highlights the specific locations within the Outeniqua Nature Reserve and the George Campus of Nelson Mandela University. Enlarged views of each location display the landscape composition and the GPS points where all scans were recorded during the study for the three baboon troops (Madiba in orange, North in dark blue, and Outeniqua in purple). Source: SANLC, 2020 - EPSG 32734. B. Principal component analysis (PCA) of land cover variables for each chacma baboon troop. The PCA plot (PC1 vs. PC2) illustrates the distribution of land cover variables with GPS points recorded every half-day for the three troops, Madiba (orange points), North (blue points), and Outeniqua (purple points).

## Materials and methods

### Study area

This study was conducted in the Western Cape, South Africa, within the Garden Route Biosphere Reserve, a 2,000 km² region extending between the cities of George and Mossel Bay (Fig. 1). The reserve, part of UNESCO’s World Network of Biosphere Reserves (Pool-Stanvliet and Coetzer 2020), includes Afro-temperate forests, fynbos shrublands, wetlands, coastal systems, and mountain ranges. The region receives a mean annual rainfall ranging from 800 to 1,100 mm, with mild temperatures typically between 18°C and 25°C (Vromans et al. 2013). Despite its ecological importance, the area has been heavily shaped by human activities, including agriculture, forestry, urban development, and tourism, especially around George, Knysna, and Mossel Bay (Socio-Economic Profile: Garden Route District Municipality, 2021). To safeguard this biodiversity hotspot, the Garden Route Biosphere Reserve was established, with the Garden Route National Park (GRNP) serving as its core, protecting vital natural areas that support biodiversity conservation. Nevertheless, the surrounding landscape forms a mosaic of cultivated, urban, and natural vegetation, producing a fragmented and patchy IUCN Category II protected area (Mormile and Hill 2017; Slater et al. 2018). Within this altered environment, numerous wildlife species, including chacma baboons, have adapted to these changing ecological conditions.

Here, three chacma baboon troops inhabiting distinct areas along a gradient of anthropogenic influence were studied to evaluate how differing degrees of human activities affect their activity patterns. The three study locations, ranging from the least to the most anthropogenically influenced, were: (i) a predominantly rural environment in Outeniqua Nature Reserve, (ii) an intermediate area between the peri-urban Campus of Nelson Mandela University and agricultural zones, and (iii) a peri-urban area within the George Campus of Nelson Mandela University. These three locations correspond hereafter to the Madiba, North, and Outeniqua troops, respectively. The Outeniqua troop (Fig. 1a, purple points), was located in the Outeniqua Nature Reserve (-33.94097, 22.42499) (Fig. 1a, purple square), covering about 38,000 ha of expansive fynbos, large tracts of native forests and pine plantations, with minimal human infrastructure (i.e. occasional buildings such as accommodation and park facilities). The North troop occupied parts of the George Campus of Nelson Mandela University (-33.96061, 22.53384) (Fig. 1a, red square) and extended its range into adjacent agricultural lands, reflecting an urban-rural transition (Fig. 1a, blue points). The campus covers approximately 400 ha, which includes a mix of natural and developed spaces, such as student housing, academic buildings, cafeterias, roads, lawns, and surrounding natural forest, fynbos, and pine plantations. The Madiba troop inhabited the same George Campus, but used more extensively built-up areas (Fig. 1a, orange points).

### Ethics

This study exclusively employed non-invasive methods, ensuring that we always maintained a minimum distance of 50 m from the chacma baboons. The research was conducted with full ethical approval from the Animal Ethics Committee of Nelson Mandela University (Permit 0544) and Cape Nature (permit CN44-87-28483). Furthermore, the study adhered to the International Primatological Society’s Code of Best Practices for Field Primatology, ensuring the highest standards of animal welfare and ethical research and the Guidelines for the Use of Animals in Behavioral Research and Teaching (Guidelines for the Use of Animals, 2012).

### Landscape characterization

To characterize the landscape used by the chacma baboons, each troop’s home range was estimated based on GPS coordinates collected during behavioural scans. For each troop, the approximate home range area was calculated during each study period using the outermost GPS locations that the group travelled during the study, using the furthest extent of their movements in all directions. Based on this method, the estimated home range sizes of the three troops were 155.9 ha for the Madiba troop, 162.3 ha for the North troop, and 164.5 ha for the Outeniqua troop (Fig. 1a). Notably, the Madiba troop had the smallest range among the three.

The study area was mapped using QGIS (3.34.11), under a WGS 84 – UTM Zone 34S projection. Landscape characterisation was made according to two environmental descriptors, the land cover and the distance to the nearest building, following the methodology described in (Kennedy Overton et al. 2024). To quantify land cover within the study area, a raster layer, obtained from the South African National Land Cover Datasets (2018), available through the Department of Forestry, Fisheries, and the Environment (DEFE, 2021) was used. The original classification included 73 distinct land cover classes, which were collapsed into 10 broader categories for analysis: natural forest, non-native forest plantations (i.e. often large pine plantation), shrubland, grassland, waterbodies, wetlands, barren land, cultivated land (i.e. any land used for growing crops, whether it’s monoculture or mixed), built-up areas, and mines. Here, built-up areas refer to all human-modified surfaces such as roads, parking lots, pavements, and other impervious infrastructure, whereas the building footprint layer specifically represents individual building structures (with roofs and walls). The building footprint layer specific to the study area was generated through geoprocessing techniques using QGIS software (Rosas-Chavoya et al. 2022). For each GPS point, land cover percentages and distance to the nearest building were calculated for each sample buffer using the Zonal Statistics plugin in QGIS. To avoid multicollinearity, variables with a correlation coefficient greater than 0.8 were excluded from the analysis. Finally, a principal component analysis (PCA), based on the habitats in which each GPS point was located, has been done to visualize the geographic distribution of the three troops and to validate the anthropogenic gradient initially established and identify potential overlap patterns.

### Baboon troop composition

The identification of individuals by age and sex followed the classification criteria outlined by (Wallace and Hill 2012). Cape chacma baboons were categorized into four groups: adult males and adult females (fully grown), subadults (individuals not fully grown, beyond juvenile in size, with unidentified sex), and juveniles (smaller than subadults, maintaining proximity to adults, with unidentified sex). The Madiba troop, the smallest of the three, included 13 individuals: 1 adult male, 3 adult females, 2 subadults, and 7 juveniles. The North troop consisted of 22 individuals, including 3 adult males, 6 adult females, 5 subadults, and 8 juveniles. Lastly, the Outeniqua troop comprised 20 individuals, including 12 adult females, 1 subadult, and 7 juveniles, with no adult male present. Across troops, Madiba, occupying the most anthropogenic environment, had the smallest home range and the smallest group size. Detailed demographic information for each of the studied troops is provided in Table 1. For this study, data collected from the Madiba, North, and Outeniqua troops are referred to in the following analysis as Madiba (M), North (N), and Outeniqua (O), respectively. New-borns <1 year were not considered or counted during this study.

**Table 1.**
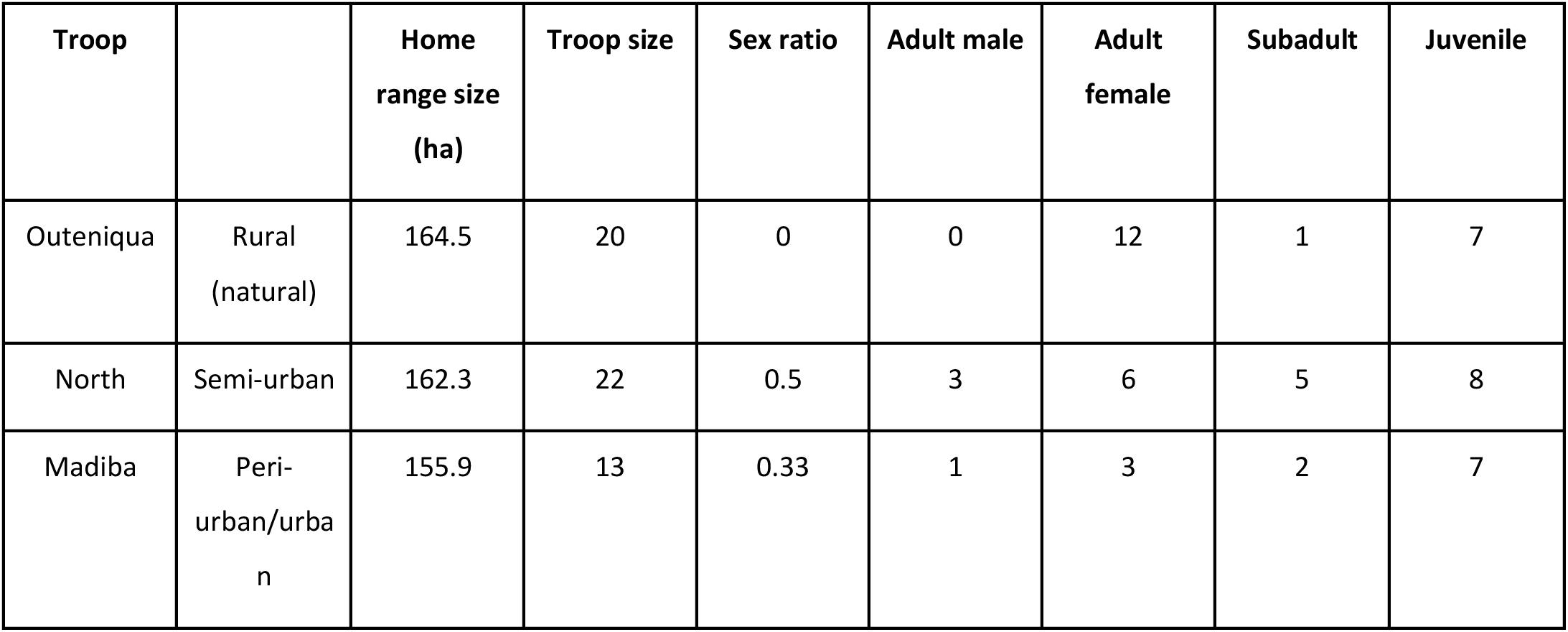
Demographic characteristics, habitat context, and home range sizes of the three chacma baboon troops included in this study. Habitat types range from rural (Outeniqua) to semi-urban (North) and peri-urban/urban (Madiba), representing a gradient of increasing anthropogenic disturbance. To characterize the landscape used by the baboons, each troop’s home range size (in hectares, ha) was estimated based on GPS coordinates recorded during behavioural scans. For each troop, the approximate home range area was calculated using the outermost GPS locations reached during the study period, representing the furthest extent of their movements in all directions. Troop composition includes all individuals observed during the study, excluding new-borns. Numbers may vary over time due to demographic events such as births, deaths, immigration, and emigration. The sex ratio is expressed as the number of adult males to adult females.

### Behavioural data collection

Data collection occurred from March 25^th^ to June 22^nd^ 2024. Observations were conducted daily between 8:00-12:00 and 13:00-17:00, with each troop observed for an average of 14 days. The minimum observation duration per half-day per troop was 5 minutes, while the maximum extended to 4 hours, depending on troop visibility and accessibility. Behavioural data were collected using scan sampling methods (Alberts and Altmann, 2006). This method was selected for its practicality under field conditions, while still producing statistically comparable results to focal sampling, the gold standard for behavioural accuracy (Gilby et al. 2010; Amato et al. 2013). The studied baboon troops were habituated to human presence. Observers maintained a standoff distance of ∼50–100 m and used video records to score behaviours without approaching or interacting with the animals.

Every 10 minutes, a scan of approximately 1.30 minutes was performed using video recording, focusing on each visible individual baboon sequentially. Video scans were performed on all visible individuals, selected in a random order to avoid sampling bias related to individual visibility or behaviour. Scans were conducted using a Canon SX70 Powershot HS camera. During each scan, we recorded: GPS coordinates (Map Marker 3.9.0_723), observer–baboon distance (Voyager laser rangefinder), habitat (open/closed), weather (sunny/cloudy), and the presence within 50 m of humans (pedestrians), other animals and cars (moving vehicles carrying humans). People inside vehicles were counted under cars, not humans, because moving vehicles represent a distinct visual and acoustic stimulus compared to pedestrians, and baboons typically respond to them differently. In addition, the spatial configuration of the group at each scan, was noted and categorized as “grouped” (all individuals within 5 meters of one another), “subgroup” (individuals within 5 meters in the same subgroup, but distant by more than 5 meters other subgroups), or “dispersed” (all individuals spaced more than 5 meters apart) (Sueur and Petit 2008).

### Behavioural video recording analysis

Following data collection, all recorded video scans were manually scored and analysed using the BORIS v.7.13.2 software program (Behavioral Observation Research Interactive Software) (Friard and Gamba 2016). The analysis was based on an ethogram specifically developed for this study, which classified observed behaviours into seven main categories: “Affiliation” (grooming, playing, lipsmacking or geckering), “Agonistic” (any form of aggressive physical contact, gesture or vocalization), “Feeding” (foraging, manipulating, or ingesting food, with the food type classified as natural or human-modified), “Locomotion” (all forms of movement, including walking, running, and fleeing), “Resting” (period of inactivity such as lying down, sitting motionless, or sleeping), “Sexual” (anogenital presentation and mounting), and “Vigilance” (distributed according to four levels of vigilance intensity, Vig_0: slow glance while in a passive position and routine scan; Vig_1: repeated glances while in a passive position and intense scanning; Vig_2: quadrupedal posture with active position and routine scanning; Vig_3: bipedal position achieving a quick glance from both sides with an active and intense stance). To enhance behavioural resolution, sub-categories were created. These included distinctions such as type of food consumed (natural vs. human-modified), mode of locomotion (walking, running, fleeing), and form of vigilance (Vigilance 0 to Vigilance 3). The complete ethogram used for this analysis is provided in Supplementary Table S1.

### Behavioural scan effort

During the study period, a total of 84 h of observation were conducted, yielding 343 behavioural scans across all three chacma baboon troops. The Madiba troop was observed for a total of 24.5 hours, during which 100 scans were recorded (61 in the morning and 39 in the afternoon) (Table 2). The North troop was monitored for 27.25 hours, with 111 behavioural scans recorded (52 morning and 59 afternoon) (Table 2). The Outeniqua troop was observed for the longest duration, 32.25 hours, with 132 scans recorded (48 in the morning and 84 in the afternoon) (Table 2).

**Table 2.** Number of behavioural scans and associated observation effort duration recorded during morning and afternoon sessions for each chacma baboon troop across the study period. Behavioural data were collected using instantaneous scan sampling every 15 minutes during two daily possible observation sessions: morning (approximately 08:00–12:00) and afternoon (approximately 13:00–17:00). The table presents the total number of scans recorded for each troop during these periods, along with the corresponding observation effort in hours. Variation in scan numbers reflects differences in troop visibility, terrain accessibility, and environmental conditions across observation days. These data were used to compute activity budgets by troop and time of day.

| Troop | Morning scans | Number of<br>morning<br>hours | Afternoon<br>scans | Number of<br>afternoon<br>hours | Total scans | Total<br>hours |
| --- | --- | --- | --- | --- | --- | --- |
| Outeniqua | 48 | 11h45 | 84 | 20h30 | 132 | 32h15 |
| North | 52 | 12h45 | 59 | 14h30 | 111 | 27h15 |
| Madiba | 61 | 15h | 39 | 9h30 | 100 | 24h30 |

### Statistical analyses

All statistical analyses were conducted using RStudio (v. 4.1.2) (Team 2024). Data normality and homoscedasticity were first assessed using the Shapiro-Wilk and the Levene’s tests (*stats* and *car* package) (Fox et al. 2001), respectively. A significance threshold of *p-value* ≤ 0.05 was used throughout. As the data did not meet the assumptions for parametric assumptions, non-parametric Kruskal-Wallis tests (*stats* package) were applied. To describe the global activity budget patterns of chacma baboons, the relative frequency of each behavioural category (Affiliation, Agonistic, Feeding, Locomotion, Resting, Sexual, and Vigilance) was calculated by dividing the frequency of each behaviour by the total number of observed behaviours for each observation period and troop (referred in the next sections to as ‘% of records’). Behavioural differences between troops (Madiba, North, and Outeniqua) were analysed using Kruskal-Wallis tests. To identify temporal variation during the day, behaviour distributions were compared between morning and afternoon periods using separate Kruskal-Wallis tests. Pairwise post-hoc comparisons between troops were performed using Dunn’s test (*FSA* package) (Ogle et al. 2015), providing *Z* values and adjusted *p*-values. Given that the Outeniqua troop was composed exclusively of adult females (Table 1), all analyses were done on two datasets: (1) the full dataset including all individuals (including females and males, hereafter referred to as the full dataset) and (2) a reduced dataset excluding males from all troops (hereafter referred to as the without-male dataset). This allowed confirmation of whether any observed patterns could be driven by troop composition or sex-specific effects. Subsequently, behavioural sub-categories nested within activity categories were analysed, which are Affiliation, Feeding, Locomotion, and Resting. These four categories were chosen because they capture the core activity budget and were the most frequent and consistently observed across troops. Agonistic, Sexual, and Vigilance behaviours were rarer, context-dependent, and could co-occur with these primary states (e.g., vigilance during feeding or resting); therefore, they were excluded from proportional activity-budget analyses to avoid bias and to maintain comparability across observation periods and groups. Kruskal-Wallis tests were used to examine how these sub-behaviours varied between troops and day periods (morning and afternoon). Post-hoc Dunn tests (*FSA* package) were again applied to identify significant pairwise differences, with corresponding *Z* and *p*-values reported.

Finally, a focused analysis of vigilance behaviour was performed, divided into four distinct categories (Vig_0 to Vig_3). The global distribution of vigilance levels across all troops, followed by a temporal comparison (morning vs. afternoon) and a troop-level comparison, was assessed using Kruskal-Wallis tests. To explore potential behavioural predictors of vigilance, we further tested whether specific vigilance categories (Vig_0 to Vig_3) were associated with three contextual variables: the individual’s position (on the ground versus in a tree), whether the behaviour occurred in a group or individual context, and the presence of external stimuli such as cars, humans, or other animals. Where relevant, post-hoc Dunn tests (*FSA* package) were applied to clarify pairwise differences. To visualize the distribution of behaviours across groups and periods, boxplots were generated using the ggplot2 package, illustrating medians, interquartile ranges, and outliers for each main behavioural category and sub-category across troop comparisons.

To assess how environmental factors influence chacma baboon activity patterns, four predictors were considered: habitat structure (open vs. non-open), weather (sunny vs. cloudy), car presence, and human presence. Habitat structure and weather were uncorrelated (χ² = 1.103, df = 1, *p*-value = 0.2936), whereas car and human presence factors were strongly correlated (Φ = 0.73). To avoid collinearity, we retained only the car variable and excluded human presence from subsequent analyses. We quantified the effects of habitat structure, weather and car presence on baboon behaviour using a series of binomial generalized linear models (GLMs) with a logit link. Five core behavioural categories were analysed (Affiliation, Feeding, Locomotion, Resting, and Vigilance) using a one-vs-all approach: for each model, scans in which the focal behaviour occurred were coded as 1 and all others as 0. Due to limited sample sizes, Agonistic (n = 23) and Sexual (n = 73) behaviours were excluded. The number of observations for each modelled category was: Affiliation (n = 1,676), Feeding (n = 2,732), Locomotion (n = 1,630), Resting (n = 1,408), and Vigilance (n = 1,940). This strategy was preferred over a multinomial model because the reduced dataset limited statistical power, while behaviour-specific binomial GLMs yielded more stable and interpretable results. In all models, habitat structure (open vs. non-open) and weather (sunny vs. cloudy) were treated as binary factors, and car presence was included as a continuous variable representing the mean number of cars observed during each scan. To account for inter-group variability, Troop identity was included as a categorical covariate in all models. Analyses were conducted twice: (1) using the full dataset (males and females combined) and (2) using a without-male subset. For the latter, the car presence variable was cleaned by converting entries to numeric and removing non-numeric values (e.g., “None”) to ensure compatibility with model fitting. For each behaviour-specific model, the Wald *p*-values associated with each environmental predictor (habitat structure, weather, car presence) were extracted. Results were summarized in dot plots, where a vertical red dashed line indicates the α = 0.05 threshold and a light grey horizontal line the *p*-value significance threshold. All analyses were implemented in R (R Core Team, 2024) using the stats package for model fitting, dplyr and tidyr for data manipulation, and ggplot2 for visualization.

## Results

### Landscape and anthropization gradient

Spatial analysis revealed marked differences in habitat use and anthropization levels among the three chacma baboon troops (Fig. 1A). The Madiba troop (orange points) was mainly located within built-up areas and surrounding shrubland and fynbos, with occasional use of adjacent natural forest patches. The North troop (blue points) showed a more balanced habitat use, occupying a mix of natural forest and grassland while still interacting with some built-up areas. In contrast, the Outeniqua troop (purple points) occurred primarily in less-disturbed environments, especially plantation forests and cultivated landscapes, and consistently avoids built-up areas. These patterns are confirmed by the PCA (Fig. 1B), which quantitatively distinguishes the spatial and environmental profiles of each troop. The PCA axis scores showed that Madiba individuals clustered closest to anthropized environments, confirming their preference for human-modified habitats (built-up). The North troop occupied an intermediate position along the anthropization gradient, reflecting mixed use of natural and semi-anthropized habitats. Finally, the Outeniqua troop was positioned furthest from anthropogenic features, indicating a strong association with natural and semi-natural environments and a preference for spatial segregation from human infrastructure. Together, these analyses confirm the differences of anthropization across the three troops, with the Madiba troop exhibiting high anthropophilic and habitat flexibility, the North troop using both human-modified and natural spaces, and the Outeniqua troop predominantly favouring more remote, less-disturbed habitats.

### Global, temporal, and troop-level activity budgets

Across the full dataset, chacma baboons allocated their time as follows: Affiliation (22.4%), Agonistic (0.4%), Feeding (35.4%), Locomotion (21.8%), Resting (19.2%), and Sexual (0.8%) (Fig. 2A). In the without-male subset, agonistic behaviour was nearly absent, indicating that adult males were the main contributors to aggressive interactions (Fig. 2B). A small number of agonistic events were nevertheless observed among Madiba females, and within this troop, adult males tended to exhibit more agonistic behaviour than females, although the difference was not statistically significant (*p*-value = 0.06781) (Fig. 2C).

**Figure 2.**
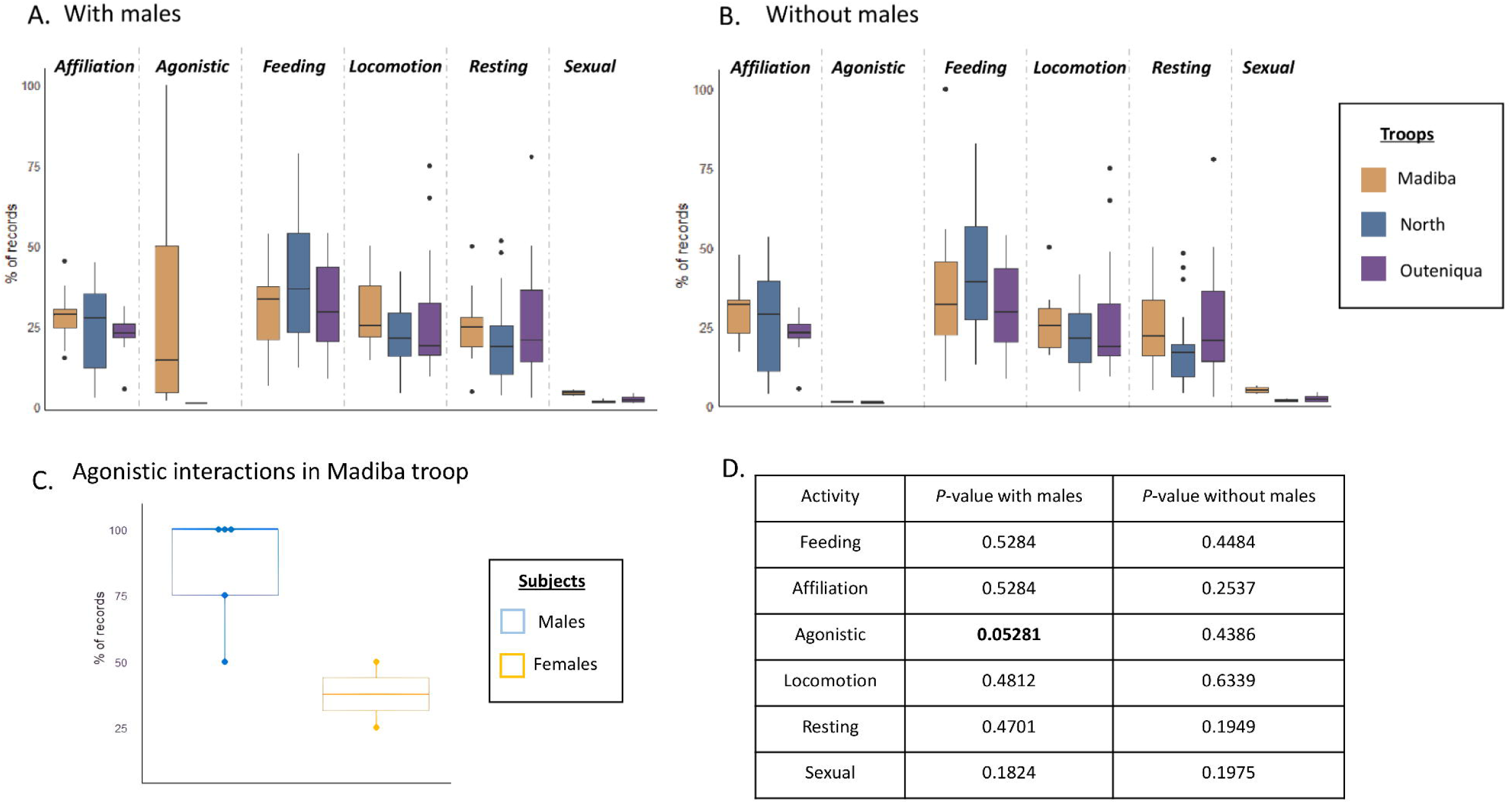
Comparison of behavioural activity patterns among chacma baboon troops and between sexes. A. Boxplots showing the percentage of scan records assigned to each of five behavioural categories (Affiliation, Feeding, Locomotion, Resting, and Vigilance) for each baboon troop: Madiba (orange), North (blue), and Outeniqua (purple), based on the full dataset (males and females). Each box represents the distribution of daily percentages of a given behaviour across scan days for each troop. The horizontal line within each box denotes the median percentage value. B. Same as panel A, but using the excluding males dataset (only females). C. Boxplots showing the proportion of Agonistic behaviour records per scan day for males and females within the Madiba troop only. Each point represents the daily proportion of Agonistic behaviour records per scan per individual sex. The horizontal line within each box represents the median proportion of Agonistic records for that sex. D. Table presenting the *p*-values for statistical comparisons of the distribution of each behavioural category between troops, for both the full dataset and the excluding males dataset (only females).

Temporally, comparing activity budgets between morning and afternoon periods revealed significant variation in several behaviours. Affiliation was significantly more frequent in the morning (24.9%) than in the afternoon (20.9%) (χ² = 5.011, df = 1, *p*-value = 0.0252), while Feeding and Locomotion were significantly more frequent in the afternoon (38.1%; 23.4%, respectively) than in the morning (30.8%; 19.0%, respectively) (χ² = 12.989, df = 1, *p*-value = 0.0003 and χ² = 6.084, df = 1, *p*-value = 0.0136, respectively). Conversely, Resting occurred more often in the morning (24.0%) than in the afternoon (16.3%) (χ² = 21.911, df = 1, *p*-value < 0.0001). There were no significant differences in the frequencies of agonistic or sexual behaviours between day periods (*p*-value > 0.90).

When assessing activity budgets by troop, normality of residuals indicated that the residuals did not follow a normal distribution (W = 0.92988, *p*-value = 1.482e-07). However, Levene’s test confirmed that the variances across groups were homogeneous (F = 0.1865, *p*-value = 0.8301), supporting the validity of statistical comparisons across troops. To further assess differences in activity budgets among the three troops, non-parametric Kruskal-Wallis tests indicated no statistically significant differences between troops for the six behavioural categories: Feeding (*p*-value = 0.5284), Affiliation (*p*-value = 0.5284), Agonistic (*p*-value = 0.0528), Locomotion (*p*-value = 0.4812), Resting (*p*-value = 0.4701), and Sexual (*p*-value = 0.1824) (Fig. 2D).

### Behavioural sub-categories

Across troops, natural food sources dominated feeding, while use of human-derived food varied. The Outeniqua troop relied significantly less on human-derived food compared to the Madiba troop (M-O: z = 2.529, *p*-value = 0.0172) (Fig. 3A). For locomotion, profiles were broadly similar with walking as the predominant mode, but Outeniqua troop used running significantly less frequently as a mode of travel compared to both the Madiba (M-O: z = 2.6904, *p*-value = 0.0107) and North troops (N-O: z = 2.5229, *p*-value = 0.0174) (Fig. 3.B). Re-analysing the data after excluding adult males yielded the same pattern, no additional contrasts became significant and effect directions were unchanged, suggesting that these troop differences are not driven by the presence of males (Fig. 3C and 3D). Overall, relative to Madiba and North, Outeniqua shows lower reliance on human foods and a lower frequency of running, against a common background of high natural-food intake and predominant walking across all troops.

**Figure 3.**
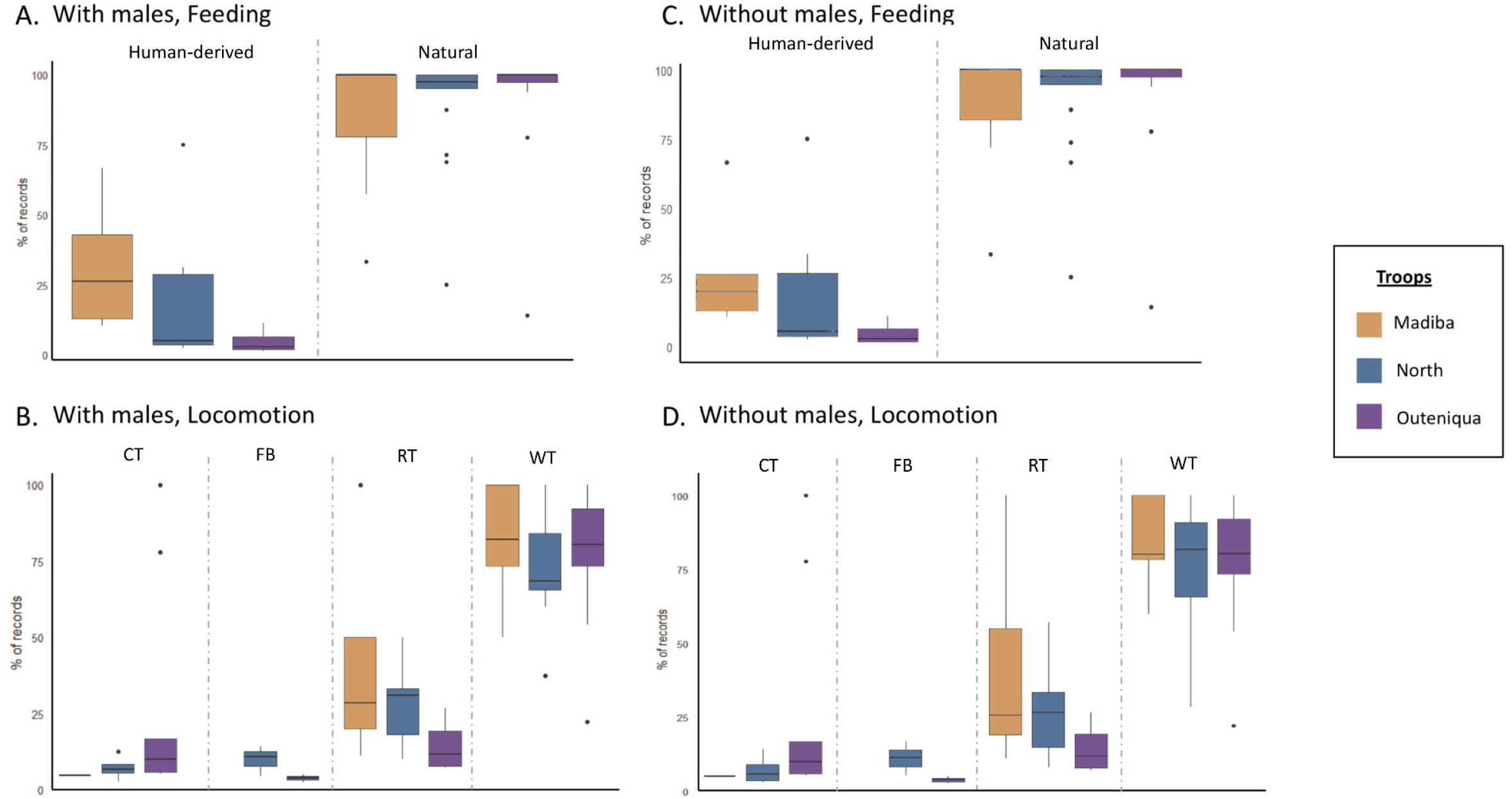
Comparison of feeding and locomotion subcategories among chacma baboon troops, with and without males. A. Boxplots showing the percentage of scan records assigned to each feeding behaviour subcategory (eating human-derived food and eating natural food) for the three baboon troops: Madiba (orange), North (blue), and Outeniqua (purple), based on the full dataset (males and females). Each box represents the distribution of daily percentages of each subcategory across scan days for each troop. The horizontal line within each box indicates the median daily proportion. B. Same as panel A, but based on the excluding males dataset (only females). C. Boxplots showing the distribution of locomotion subcategories across the three troops using the full dataset. Locomotion subcategories include: CT (Climbing Travelling), FB (Flight Baboon), RT (Running Travelling), and WT (Walking Travelling). Each box summarizes the daily percentage of records attributed to each locomotion subcategory, with the median indicated by the horizontal line. D. Same as panel C, but using the excluding males dataset (only females).

### Environmental drivers of activity

In the full dataset (males + females; Fig. 4A; Supplementary Table 2), Feeding increased significantly in open habitats (β = 0.60 ± 0.15, *p*-value = 0.00012) and under sunny weather (β = 0.26 ± 0.09, *p*-value = 0.0035) but decreased with higher car presence (β = −0.16 ± 0.03, *p*-value < 0.0001). Vigilance showed an opposite trend, with significantly lower occurrence in sunny conditions (β = −0.18 ± 0.07, *p*-value = 0.015) and a marked increase with car presence (β = 0.24 ± 0.03, *p*-value < 10⁻¹⁷). Locomotion declined significantly in open habitats (β = −0.36 ± 0.15, *p*-value = 0.017) and with greater car presence (β = −0.20 ± 0.04, *p*-value < 10⁻⁶). Resting increased under sunny conditions (β = 0.40 ± 0.12, *p*-value = 0.0008) but declined with car presence (β = −0.10 ± 0.04, *p*-value = 0.015). Affiliation was less frequent in open habitats (β = −0.37 ± 0.15, *p*-value = 0.014) and under sunny weather (β = −0.38 ± 0.10, *p*-value = 0.0001), suggesting that social interactions were reduced in exposed conditions.

**Figure 4.**
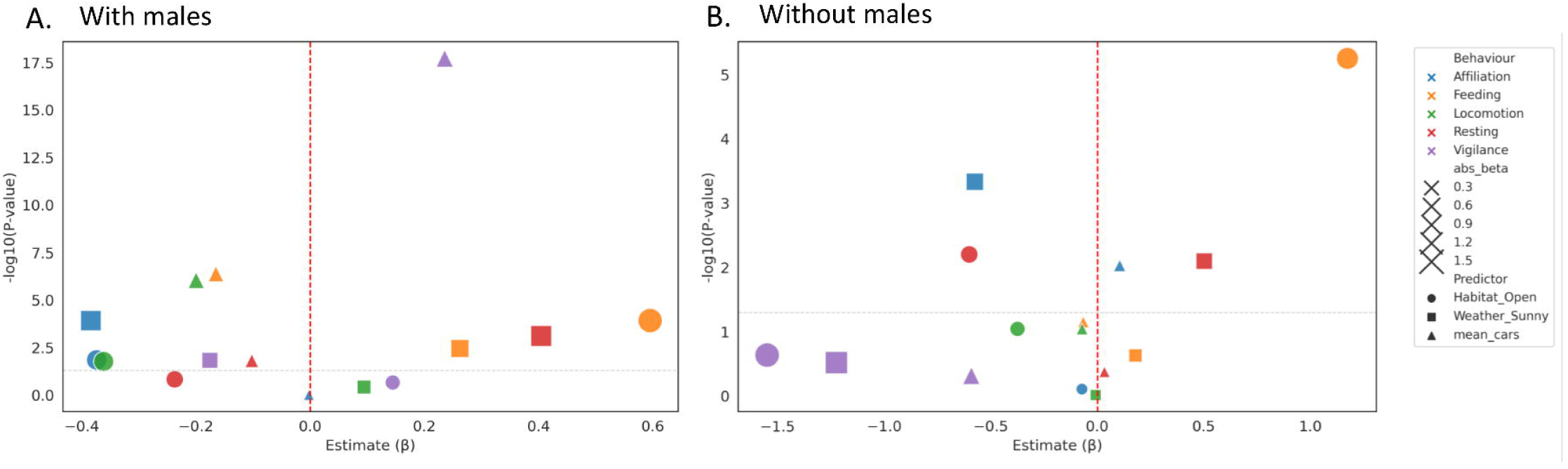
Effects of environmental predictors on chacma baboon behaviour. Bubble plots show the direction, magnitude, and statistical significance of environmental predictors on five behavioural categories (Affiliation in blue, Feeding in orange, Locomotion in green, Resting in red, and Vigilance in purple) derived from binomial generalized linear models (GLMs). Each point represents a predictor–behaviour combination. The x-axis shows the estimated regression coefficient (β), with positive values indicating an increase and negative values a decrease in the probability of a given behaviour. The y-axis shows significance as –log₁₀(*p*-value); the light grey dashed line corresponds to the *p*-value = 0.05 threshold (–log₁₀ ≈ 1.3), and the vertical red dashed line marks β = 0. Point shape indicates the environmental predictor (● Habitat openness, ▪ Weather, ▴ Mean number of cars), and bubble size is proportional to the absolute effect magnitude (|β|). A. Effects of environmental predictors for the full dataset including males and females. B. Effects of environmental predictors for the male excluding dataset. Corresponding model coefficients (β), standard errors (SE), Wald z-values, and *p*-values are reported in Tables S2 and S3.

When only females were considered (i.e. without-male dataset, Fig. 4B; Supplementary Table 3), overall model significance decreased, but some consistent trends persisted. Feeding remained strongly associated with open habitats (β = 1.17 ± 0.26, *p*-value < 0.00001), confirming that this behavioural response is robust to troop sex composition. Resting increased in sunny weather (β = 0.50 ± 0.19, *p*-value = 0.008) but decreased in open habitats (β = −0.60 ± 0.22, *p*-value = 0.006). Affiliation was negatively affected by sunny weather (β = −0.58 ± 0.16, *p*-value = 0.0005) but showed a slight positive association with car presence (β = 0.11 ± 0.04, *p*-value = 0.009), suggesting that females may increase social interactions under moderate anthropic disturbance. Other behaviours (Locomotion and Vigilance) exhibited non-significant effects (*p*-value > 0.05), suggesting weaker environmental sensitivity.

These findings demonstrate that weather, habitat openness, and human disturbance interact to baboon activity patterns, influencing how individuals allocate time between vigilance, movement, and foraging across varying environmental contexts (see Supplementary Tables S2 and S3 for full GLM outputs).

### Vigilance patterns

Across all vigilance observations, chacma baboons expressed the following overall proportions of vigilance: vigilance 0 (Vig_0; no vigilance; 79.1% of all scan records), vigilance 1 (Vig_1; occasional scanning; 8.4%), vigilance 2 (Vig_2; regular scanning; 7.6%) and vigilance 3 (Vig_3; intense, active scanning; 4.9%) (Fig. 5A). When comparing vigilance among troops, only the third level of vigilance (Vig_3) differed significantly across groups (*p*-value = 0.021). The Outeniqua troop exhibited this high-intensity vigilance least frequently (Madiba–Outeniqua: z = 3.40, *p*-value = 0.0010; North–Outeniqua: z = 2.35, *p*-value = 0.0279) (Fig. 5B). Mean vigilance 3 proportions per troop were Madiba: 7.8 ± 2.9%, North: 6.3 ± 2.5%, and Outeniqua: 2.1 ± 1.0% of total scans. When adult males were excluded from the dataset, no significant between-troop differences were detected at any vigilance level (all pairwise contrasts *p*-value > 0.05; Fig. 5C–D), suggesting that the previously observed difference in vigilance 3 is primarily driven by adult male behaviour.

**Figure 5.**
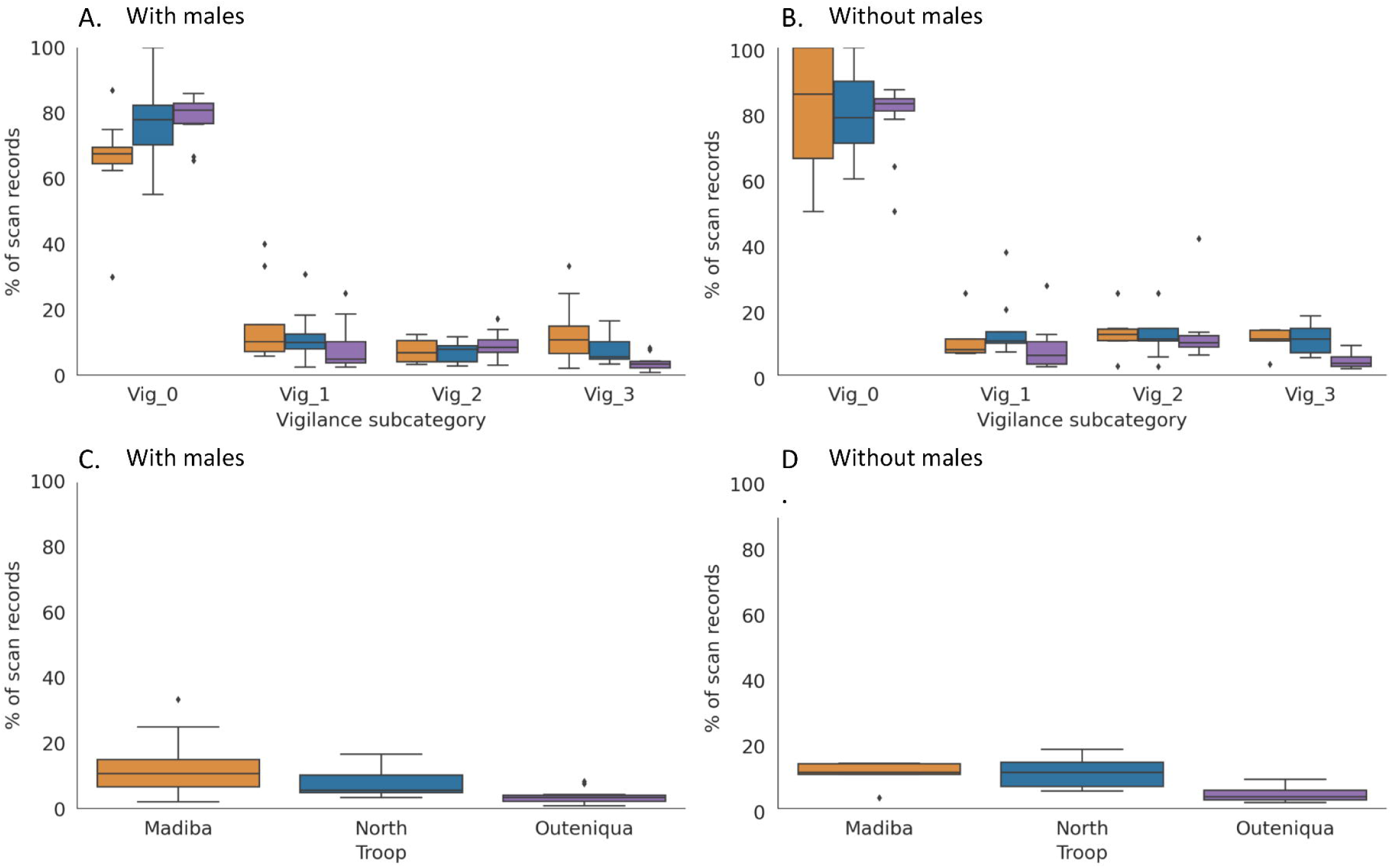
Comparison of vigilance among chacma baboon troops, with and without males. A. Boxplots showing the percentage of scan records assigned to each vigilance subcategory (Vig_0, Vig_1, Vig_2 and Vig_3) for the three baboon troops: Madiba (orange), North (blue), and Outeniqua (purple), based on the full dataset (males and females). Each box represents the distribution of daily percentages of each subcategory across scan days for each troop. The horizontal line within each box indicates the median daily proportion. B. Same as panel A, but based on the excluding males dataset (female-only). C. Boxplots showing the distribution of the vigilance 3 across the three troops using the full dataset. Each box summarizes the daily percentage of records attributed to the vigilance 3 subcategory only, with the median indicated by the horizontal line. D. Same as panel C, but using the excluding males dataset.

## Discussion

Anthropogenic landscapes restructure resource availability and risk, prompting predictable shifts in primate time budgets and vigilance. The main aim of the present study was to evaluate how a rural–peri-urban gradient and associated anthropogenic factors reshape chacma baboon activity budgets and vigilance. Controlling for weather, habitat structure, car and human presence, the peri-urban troop (Madiba) showed clear signatures of anthropogenic foraging (greater use of human-derived foods) and distinct locomotion relative to the rural context, without clear evidence for a wholesale reallocation toward low-cost behaviours such as resting; instead, disturbance (car/human presence) was associated with context-dependent increases in vigilance and locomotion (H1). As anticipated, the intermediate troop (North) generally occupied an intermediate position across behavioural metrics, though not uniformly so (H2). With respect to vigilance, only the highest-intensity state (Vig_3) differed among troops in the full dataset, with the Outeniqua troop inhabiting the most rural area showing the lowest rate; this difference vanished when adult males were excluded, indicating that between-troop variation in intense scanning is largely male-driven (H3). Together, these results suggest that variation among troops arises from a complex combination of resource availability (access to anthropogenic foods) and disturbance-mediated risk responses, modulated by sex composition rather than anthropogenic setting alone.

### Land uses and anthropogenic profiling

In this study, quantitative land-use analyses confirmed contrasts in habitat use among the three troops. (Fig. 1A–B). Outeniqua troop primarily used natural habitats (e.g., plantation forest, cultivated fields) and seemed to avoid built-up areas. North troop displayed an intermediate profile, splitting use between natural habitats (grasslands, forests) and occasional incursions into built-up areas, similar to patterns reported for olive baboons in mixed agro-natural mosaics (Gilbert 2017). Madiba troop showed the strongest association with human-modified areas, frequently using built-up zones and adjacent shrubland, which is consistent with prior work identifying this troop as the most anthropophilic via analysis of faeces stable isotopes showing a higher ratio of nitrogen isotope, a pattern typically associated to the ingestion of human-made food (Kennedy Overton et al. 2024) and other studies on Cape Town chacma baboons (Hoffman and O’Riain 2012). Together, these patterns align troops flexibly exploitation depending on the ecological opportunities and anthropogenic disturbance (Hoffman and O’Riain 2012).

### Anthropogenic environments impact on food type, locomotion and vigilance

Across all studied troops, feeding dominated the daily activity budget (35.4%), matching already known classic patterns in the genus *Papio* and other primates where food acquisition is the central time sink (Altmann and Muruthi 1988; Barrett et al. 2019). In the studied dataset, feeding was followed by non-feeding activities, resting, moving, affiliation, and vigilance, in the expected proportions for Cape chacma baboons inhabiting heterogeneous mosaic areas. This rank order is consistent with broader primate literature showing high feeding investment under patchy or seasonal resources (e.g., chimpanzees and colobus) (Goodall 1986; Oates 1994; Balcomb et al. 2000). A notable nuance concerns agonistic behaviour, which approached significance in the most urban inhabited area when adult males were included. This pattern may likely reflect male-specific competition related to mating opportunities, dominance status, coalition formation, territorial defence, or mate-guarding (Colmenares 1991; Patzelt et al. 2014; Cerda-Molina et al. 2023). Together, these results situate Cape chacma baboons from the Garden Route within the expected primate activity framework while highlighting how sex composition may shape the expression of aggression.

No overall differences were detected in total activity budgets among the three troops. However, behaviour-specific shifts are partly consistent with H1 hypothesis, with contrasts emerging within specific behavioural sub-categories, with Outeniqua differing significantly from Madiba and/or North troops, depending on the behaviour considered. Regarding the food type, the Outeniqua troop relied significantly less on human-derived food compared to the Madiba troop (p = 0.0172), consistent with its lower access to anthropogenic resources and with studies showing that primates in less disturbed habitats depend more on natural foods (e.g., McLennan and Hill, 2012). A study by McLennan and Hill showed that chimpanzee communities in less disturbed areas in Uganda consumed a higher proportion of wild fruits and leaves compared to those in areas with greater human presence, where they resorted to crop-foraging (McLennan and Hill 2012). We did not find any difference in natural food consumption between the troops, likely because the campus landscape still contains substantial natural formations (i.e. native forest, shrubland and fynbos; Fig. 1.A-B), providing reliable fall-back resources when human-derived foods are scarce. The study by Mazué and al. carried out on the same two troops, Madiba and North respectively, confirmed that baboon troops were very good at adjusting their food sources according to the availability of human food resources, and were therefore able when necessary to focus on foraging in the natural environment (Mazué et al. 2023). In terms of locomotion, the Outeniqua troop ran significantly less than both the Madiba (*p*-value = 0.0107) and North troops (*p*-value = 0.0174). Similar patterns have been reported across primates, with locomotor shifts tracking habitat structure and resource distribution. Although spider monkeys (*Ateles* spp.) are predominantly arboreal, occasional increases in terrestrial locomotion have been interpreted as escape/avoidance responses under disturbance rather than routine travel per se (Fleagle 2015). In our system, the elevated running we observed in more anthropogenic settings is therefore more parsimoniously explained by a disturbance-related function (e.g., rapid displacement away from humans or vehicles) than by openness alone. These results align partially with H2, with the North troop generally occupying an intermediate position between Outeniqua and Madiba, and differing significantly from Outeniqua (e.g., running, *p* = 0.0174), but it is not consistently distinct from Madiba, so a strict ordering (Outeniqua < North < Madiba) is not always resolved. Overall, the patterns observed fit broader primate responses to anthropogenic environments, with variation in reliance on human-derived foods and locomotion adjustments underscoring the behavioural flexibility of chacma baboons in the face of shifting ecological, social and anthropogenic pressures.

Vigilance behaviour appears different between troops, with the Outeniqua troop (the troop living in the less anthropogenic environment) showing a significantly lower third level of vigilance (Vig_3; active posture and intense vigilance) than the Madiba and the North troops, which are more often in contact with human activities and infrastructures, partially supporting H2. This mirrors previous research showing that primates’ vigilance is highly influenced by environmental context. For instance, a study on Toque macaques (*Macaca sinica*) and one on rhesus macaques (*Macaca mulatta*) found that the time devoted to vigilance was significantly higher and more present in “moderately to highly urbanized” sites, compared to the “weakly urbanized” site, similar to the heightened vigilance observed in the Madiba and North troops (Kaburu et al. 2019; Jayapali et al. 2023). Together, these results underscore that along the rural-urban gradient, vigilance is the most sensitive behavioural axis, with diet-source use and running varying in parallel, even in well-habituated groups.

### Habitat structure and car exposure as main drivers of behavioural flexibility

Environmental context is a major determinant of primate behavioural flexibility (Kappeler et al. 2013; Schradin 2013). In this study, habitat structure (openness vs. closure) and an anthropogenic disturbance (presence of cars) emerged as the most influential predictors shaping baboon activity. In the full dataset, feeding and vigilance were the behaviours most responsive to environmental variation. Feeding increased markedly in open habitats and under sunny conditions, while vigilance decreased in similar contexts but increased with car presence, indicating contrasting behavioural strategies under changing environmental conditions. These results suggest that open and more disturbed environments promote greater foraging activity, possibly due to enhanced visibility and accessibility of food, but at the cost of reduced vigilance, a trade-off between foraging efficiency and risk assessment (Bersacola et al. 2021; Hammond et al. 2022). Habitat openness also affected social behaviours, with affiliative interactions occurring less frequently in open habitats (*p*-value = 0.014), consistent with the idea that reduced refuge availability and higher exposure constrain time for close social contact. Bolt and al. showed that Geoffroy’s spider monkeys (*Ateles geoffroyi*), Panamian white-throated capuchin (*Cebus imitator*) and Mantled howler monkeys (*Alouatta palliata*) express lower rates and altered types of social behaviour in the forest fringe than in the forest interior, reducing or modifying their affiliative social behaviours according to their habitat structure (Bolt et al. 2025).

The presence of cars, used as a proxy for human disturbance, exerted a more behaviour-specific influence. In the full dataset, resting decreased with higher car density (*p*-value = 0.015), indicating that anthropogenic disturbance limits opportunities for rest. However, in the without-male dataset, this relationship weakened, and feeding responses to habitat openness remained the only robust pattern (*p*-value < 0.001), suggesting that males may drive part of the behavioural variance in vigilance and resting patterns. Interestingly, affiliation in females was slightly positively associated with car presence (*p*-value = 0.009), possibly reflecting compensatory social bonding under moderate disturbance. Several studies on long-tailed macaques (*Macaca fascicularis*), olive baboons (*Papio anubis*), or vervet monkeys *(Chlorocebus aethiops)* have also highlighted adjustments in their activities in urban settings, particularly a reduction in the time available for social interactions, grooming, resting, and modifications in their eating habits (Strum 2010; Sha and Hanya 2013; Novak et al. 2014; Thatcher et al. 2020). However, this variable did not influence the vigilance of the baboons, reflecting a possible habituation of the species, which no longer considers these stimuli as a threat. Finally, through this analysis, activity patterns of baboons seem to be driven by an association of natural environmental conditions and anthropogenic environmental conditions.

## Conclusion

Across this rural-urban gradient, Cape chacma baboons showed canonical time allocation (feeding dominant), with no whole-budget differences among troops but fine behaviour-specific shifts with greater use of human-derived foods and more running in the more anthropogenic contexts, and vigilance as the most sensitive behaviour (Madiba/North > Outeniqua). H1 hypothesis was partially supported at the behavioural level (not at the whole budget), H2 was also partially supported (higher vigilance in peri-urban Madiba vs Outeniqua), and the H3 (North intermediate) was directionally supported but not always statistically distinct. Environmental covariates, habitat openness (lower affiliation) and traffic (lower resting, no effect on vigilance), help to better understand these patterns.

Some key limitations need to be highlighted in the present study such as the lack of within-environment population replicates (potential troop-identity effects), possible seasonal confounding (despite weather terms, longer time series would improve inference), and unmeasured fine-scale variation in human food availability and interaction frequency. Moreover, observations over a longer period (with a greater number of observation hours) would help refine our results. These are partly mitigated by explicit land-use profiling, concordance with prior studies, and the covariate results, but they warrant caution. In such a context, in the future studies should add replicate troops per environmental category (a quite complicated approach to implement), follow multi-season/annual sampling, and add other approaches such as fine-scale GPS/accelerometers data with concurrent measures of food access, traffic, and people presence. Experimental or quasi-experimental management studies (securing anthropogenic food, traffic calming at crossings) could help testing causality and, if effective, reduce vigilance and conflict while preserving natural foraging opportunities. Such approaches will help resolving how anthropogenic disturbance reshapes primate time allocation and risk perception, which is essential for evidence-based conservation and conflict mitigation.

## Credit authorship contribution statement

Soanna Dany: Conceptualization, Investigation, Methodology, Field work, Data curation, Formal analysis, Writing – original draft, review & editing. Celia Lacomme: Methodology, Writing – review & editing. Anne Guionneau: Field work, Writing – review & editing. Aluwani Ramaru: Field work. Franck Prugnolle: Supervision, writing – review & editing, Odile Petit: Conceptualization, Supervision, Investigation, Methodology, Data curation, Formal analysis, Writing – review & editing, Virginie Rougeron: Conceptualization, Supervision, Investigation, Methodology, Data curation, Formal analysis, Writing – original draft, review & editing, Funding acquisition.

## Acknowledgements

We are grateful to SANParks for allowing and participating in data collection throughout this study. We would especially like to thank all involved assistants who provided us access and guidance throughout the study areas. Thank you to all the people who provided details on troop locations throughout the Garden Route.

## References

Cowlishaw G & Robin I. M. Dunbar. 2000. Primate Conservation Biology. 1st editio. University of Chicago Press.

Coppolillo MBM & P. 2005. Conservation: Linking Ecology, Economics, and Culture. Princeton University Press.

Estrada A et al. 2017. Impending extinction crisis of the world’s primates: Why primates matter. Sci Adv. 3(1) https://www.science.org/doi/10.1126/sciadv.1600946. 10.1126/sciadv.1600946

Vitousek PM, Mooney HA, Lubchenco J, Melillo JM. 1997. Human Domination of Earth’s Ecosystems. Science (80). 277(5325):494–499 https://www.science.org/doi/10.1126/science.277.5325.494. 10.1126/science.277.5325.494

Tilman D et al. 2017. Future threats to biodiversity and pathways to their prevention. Nature. 546(7656):73–81 https://www.nature.com/articles/nature22900. 10.1038/nature22900

Achard F, Eva H, Mayaux P. 2001. Tropical forest mapping from coarse spatial resolution satellite data: Production and accuracy assessment issues. Int J Remote Sens. 22(14):2741–2762 https://www.tandfonline.com/doi/full/10.1080/01431160120548. 10.1080/01431160120548

Muche M et al. 2023. Land use and land cover changes and their impact on ecosystem service values in the north-eastern highlands of Ethiopia Veettil BK, editor. PLoS One. 18(9):e0289962 https://dx.plos.org/10.1371/journal.pone.0289962. 10.1371/journal.pone.0289962

Hulme-Beaman A, Dobney K, Cucchi T, Searle JB. 2016. An Ecological and Evolutionary Framework for Commensalism in Anthropogenic Environments. Trends Ecol Evol. 31(8):633–645 https://linkinghub.elsevier.com/retrieve/pii/S0169534716300441. 10.1016/j.tree.2016.05.001

La Deau S, Leisnham P, Biehler D, Bodner D. 2013. Higher Mosquito Production in Low-Income Neighborhoods of Baltimore and Washington, DC: Understanding Ecological Drivers and Mosquito-Borne Disease Risk in Temperate Cities. Int J Environ Res Public Health. 10(4):1505–1526 https://www.mdpi.com/1660-4601/10/4/1505. 10.3390/ijerph10041505

Murray MH, St. Clair CC. 2017. Predictable features attract urban coyotes to residential yards. J Wildl Manage. 81(4):593–600 https://wildlife.onlinelibrary.wiley.com/doi/10.1002/jwmg.21223. 10.1002/jwmg.21223

Prange, S. and Gehrt, S. D. and Wiggers EP. 2004. Influences of Anthropogenic Resources on Raccoon (*Procyon lotor*) Movements and Spatial Distribution. J Mammal. [published online ahead of print] https://academic.oup.com/jmammal/article/85/3/483/901013/Influences-of-Anthropogenic-Resources-on-Raccoon. 10.1644/1383946

Dugger KM et al. 2016. The effects of habitat, climate, and Barred Owls on long-term demography of Northern Spotted Owls. Condor. 118(1):57–116 https://academic.oup.com/condor/article/118/1/57/5153234. 10.1650/CONDOR-15-24.1

Maisels F et al. 2013. Devastating Decline of Forest Elephants in Central Africa Kolokotronis S-O, editor. PLoS One. 8(3):e59469 https://dx.plos.org/10.1371/journal.pone.0059469. 10.1371/journal.pone.0059469

Tempel DJ et al. 2014. Effects of forest management on California Spotted Owls: implications for reducing wildfire risk in fire-prone forests. Ecol Appl. 24(8):2089–2106 https://esajournals.onlinelibrary.wiley.com/doi/10.1890/13-2192.1. 10.1890/13-2192.1

Larson L, Conway A, Hernandez S, Carroll J. 2016. Human-wildlife Conflict, Conservation Attitudes, and a Potential Role for Citizen Science in Sierra Leone, Africa. Conserv Soc. 14(3):205 https://journals.lww.com/10.4103/0972-4923.191159. 10.4103/0972-4923.191159

Schell CJ et al. 2021. The evolutionary consequences of human–wildlife conflict in cities. Evol Appl. 14(1):178–197 https://onlinelibrary.wiley.com/doi/10.1111/eva.13131. 10.1111/eva.13131

Bateman PW, Fleming PA. 2012. Big city life: carnivores in urban environments Le Comber S, editor. J Zool. 287(1):1–23 https://zslpublications.onlinelibrary.wiley.com/doi/10.1111/j.1469-7998.2011.00887.x. 10.1111/j.1469-7998.2011.00887.x

Hetmański T, Bocheński M, Tryjanowski P, Skórka P. 2011. The effect of habitat and number of inhabitants on the population sizes of feral pigeons around towns in northern Poland. Eur J Wildl Res. 57(3):421–428 http://link.springer.com/10.1007/s10344-010-0448-z. 10.1007/s10344-010-0448-z

Scholz C et al. 2020. Individual dietary specialization in a generalist predator: A stable isotope analysis of urban and rural red foxes. Ecol Evol. 10(16):8855–8870 https://onlinelibrary.wiley.com/doi/10.1002/ece3.6584. 10.1002/ece3.6584

Charmantier A, Gienapp P. 2014. Climate change and timing of avian breeding and migration: evolutionary versus plastic changes. Evol Appl. 7(1):15–28 https://onlinelibrary.wiley.com/doi/10.1111/eva.12126. 10.1111/eva.12126

Sih A, Trimmer PC, Ehlman SM. 2016. A conceptual framework for understanding behavioral responses to HIREC. Curr Opin Behav Sci. 12:109–114 https://linkinghub.elsevier.com/retrieve/pii/S2352154616301887. 10.1016/j.cobeha.2016.09.014

Bersacola E et al. 2023. Primate Conservation in Shared Landscapes. p 161–181 https://link.springer.com/10.1007/978-3-031-11736-7_10. 10.1007/978-3-031-11736-7_10

McLennan MR, Spagnoletti N, Hockings KJ. 2017. The Implications of Primate Behavioral Flexibility for Sustainable Human–Primate Coexistence in Anthropogenic Habitats. Int J Primatol. 38(2):105–121 http://link.springer.com/10.1007/s10764-017-9962-0. 10.1007/s10764-017-9962-0

McLennan MR, Lorenti GA, Sabiiti T, Bardi M. 2020. Forest fragments become farmland: Dietary Response of wild chimpanzees (*Pan troglodytes*) to fast-changing anthropogenic landscapes. Am J Primatol. 82(4) https://onlinelibrary.wiley.com/doi/10.1002/ajp.23090. 10.1002/ajp.23090

Corrêa FM, Chaves ÓM, Printes RC, Romanowski HP. 2018. Surviving in the urban–rural interface: Feeding and ranging behavior of brown howlers (Alouatta guariba clamitans) in an urban fragment in southern Brazil. Am J Primatol. 80(6) https://onlinelibrary.wiley.com/doi/10.1002/ajp.22865. 10.1002/ajp.22865

Dhananjaya T et al. 2022. Extent of encounter with an embedded food influences how it is processed by an urbanizing macaque species. Behaviour. 159(7):657–689 https://brill.com/view/journals/beh/159/7/article-p657_4.xml. 10.1163/1568539X-bja10146

Fehlmann G et al. 2017. Extreme behavioural shifts by baboons exploiting risky, resource-rich, human-modified environments. Sci Rep. 7(1):15057 https://www.nature.com/articles/s41598-017-14871-2. 10.1038/s41598-017-14871-2

Kaplan BS, O’Riain MJ, van Eeden R, King AJ. 2011. A Low-Cost Manipulation of Food Resources Reduces Spatial Overlap Between Baboons (*Papio ursinus*) and Humans in Conflict. Int J Primatol. 32(6):1397–1412 http://link.springer.com/10.1007/s10764-011-9541-8. 10.1007/s10764-011-9541-8

Katlam G, Prasad S, Aggarwal M, Kumar R. 2018. Trash on the Menu:Patterns of Animal Visitation and Foraging Behaviour at Garbage Dumps. Curr Sci. 115(12):2322 https://www.currentscience.ac.in/Volumes/115/12/2322.pdf. 10.18520/cs/v115/i12/2322-2326

Kennedy Overton E et al. 2024. Land use influences the diet of chacma baboons (Papio ursinus) in South Africa. Glob Ecol Conserv. 54:e03100 https://linkinghub.elsevier.com/retrieve/pii/S2351989424003044. 10.1016/j.gecco.2024.e03100

Saj TL, Sicotte P, Paterson JD. 2001. The conflict between vervet monkeys and farmers at the forest edge in Entebbe, Uganda. Afr J Ecol. 39(2):195–199 https://onlinelibrary.wiley.com/doi/10.1046/j.0141-6707.2000.00299.x. 10.1046/j.0141-6707.2000.00299.x

Hoffman TS, O’Riain MJ. 2012. Landscape requirements of a primate population in a human-dominated environment. Front Zool. 9(1):1 http://frontiersinzoology.biomedcentral.com/articles/10.1186/1742-9994-9-1. 10.1186/1742-9994-9-1

Oi T, Hamasaki S, Seino H, Kawamoto Y. 2021. Inter-individual variation in the diet within a group of Japanese macaques and its relationship with social structure investigated by stable isotope and DNA analyses. Primates. 62(1):103–112 https://link.springer.com/10.1007/s10329-020-00840-3. 10.1007/s10329-020-00840-3

Riley EP, Fuentes A. 2011. Conserving social–ecological systems in Indonesia: human–nonhuman primate interconnections in Bali and Sulawesi. Am J Primatol. 73(1):62–74 https://onlinelibrary.wiley.com/doi/10.1002/ajp.20834. 10.1002/ajp.20834

Novak MA, El-Mallah SN, Menard MT. 2014. Use of the Cross-Translational Model to Study Self-Injurious Behavior in Human and Nonhuman Primates. ILAR J. 55(2):274–283 https://academic.oup.com/ilarjournal/article-lookup/doi/10.1093/ilar/ilu001. 10.1093/ilar/ilu001

Kumara HN, Kumar S, Singh M. 2010. Of how much concern are the ‘least concern’ species? Distribution and conservation status of bonnet macaques, rhesus macaques and Hanuman langurs in Karnataka, India. Primates. 51(1):37–42 http://link.springer.com/10.1007/s10329-009-0168-8. 10.1007/s10329-009-0168-8

Lewis MC, O’Riain MJ. 2017. Foraging Profile, Activity Budget and Spatial Ecology of Exclusively Natural-Foraging Chacma Baboons (*Papio ursinus*) on the Cape Peninsula, South Africa. Int J Primatol. 38(4):751–779 http://link.springer.com/10.1007/s10764-017-9978-5. 10.1007/s10764-017-9978-5

Sha JCM, Hanya G. 2013. Diet, Activity, Habitat Use, and Ranging of Two Neighboring Groups of Food-Enhanced Long-Tailed Macaques (*Macaca fascicularis*). Am J Primatol. 75(6):581–592 https://onlinelibrary.wiley.com/doi/10.1002/ajp.22137. 10.1002/ajp.22137

Thatcher HR, Downs CT, Koyama NF. 2020. Understanding foraging flexibility in urban vervet monkeys, Chlorocebus pygerythrus, for the benefit of human-wildlife coexistence. Urban Ecosyst. 23(6):1349–1357 https://link.springer.com/10.1007/s11252-020-01014-1. 10.1007/s11252-020-01014-1

Almeida-Rocha JM de, Peres CA, Oliveira LC. 2017. Primate responses to anthropogenic habitat disturbance: A pantropical meta-analysis. Biol Conserv. 215:30–38 https://linkinghub.elsevier.com/retrieve/pii/S000632071630893X. 10.1016/j.biocon.2017.08.018

Henzi P, Barrett L. 2003. Evolutionary ecology, sexual conflict, and behavioral differentiation among baboon populations. Evol Anthropol Issues, News, Rev. 12(5):217–230 https://onlinelibrary.wiley.com/doi/10.1002/evan.10121. 10.1002/evan.10121

Alberts SC, Altmann J. 2006. The Evolutionary Past and the Research Future: Environmental Variation and Life History Flexibility in a Primate Lineage. In: Reproduction and fitness in baboons: behavioral, ecological, and life history perspectives. Springer US; p 277–303 http://link.springer.com/10.1007/0-387-33674-5_12. 10.1007/0-387-33674-5_12

Else JG. 1991. Non human primates as pests. H. O. Box. London: Chapman & Hall.; p 155–166

Whiten A, Byrne RW, Henzi SP. 1987. The behavioral ecology of mountain baboons. Int J Primatol. 8(4):367–388 http://link.springer.com/10.1007/BF02737389. 10.1007/BF02737389

Whiten, A. and Byrne, R. W. and Barton, R. A. and Waterman, P. G. and Henzi SP. 1991. Dietary and foraging strategies of baboons. Philos Trans R Soc London Ser B Biol Sci. 334(1270):187–197 https://royalsocietypublishing.org/doi/10.1098/rstb.1991.0108. 10.1098/rstb.1991.0108

Chowdhury S, Brown J, Swedell L. 2020. Anthropogenic effects on the physiology and behaviour of chacma baboons in the Cape Peninsula of South Africa Cooke S, editor. Conserv Physiol. 8(1) https://academic.oup.com/conphys/article/doi/10.1093/conphys/coaa066/5879264. 10.1093/conphys/coaa066

Hill CM. 2000. Conflict of Interest Between People and Baboons: Crop Raiding in Uganda. Int J Primatol. 21(2):299–315 https://link.springer.com/10.1023/A:1005481605637. 10.1023/A:1005481605637

Beamish EK, O’Riain MJ. 2014. The Effects of Permanent Injury on the Behavior and Diet of Commensal Chacma Baboons (*Papio ursinus*) in the Cape Peninsula, South Africa. Int J Primatol. 35(5):1004–1020 http://link.springer.com/10.1007/s10764-014-9779-z. 10.1007/s10764-014-9779-z

Henzi SP, Brown LR, Barrett L, Marais AJ. 2011. Troop Size, Habitat Use, and Diet of Chacma Baboons (*Papio hamadryas ursinus*) in Commercial Pine Plantations: Implications for Management. Int J Primatol. 32(4):1020–1032 http://link.springer.com/10.1007/s10764-011-9519-6. 10.1007/s10764-011-9519-6

Mazué F et al. 2023. Less bins, less baboons: reducing access to anthropogenic food effectively decreases the urban foraging behavior of a troop of chacma baboons (*Papio hamadryas ursinus*) in a peri-urban area. Primates. 64(1):91–103 https://link.springer.com/10.1007/s10329-022-01032-x. 10.1007/s10329-022-01032-x

Pebsworth PA, MacIntosh AJJ, Morgan HR, Huffman MA. 2012. Factors Influencing the Ranging Behavior of Chacma Baboons (*Papio hamadryas ursinus*) Living in a Human-Modified Habitat. Int J Primatol. 33(4):872–887 http://link.springer.com/10.1007/s10764-012-9620-5. 10.1007/s10764-012-9620-5

Pool-Stanvliet R, Coetzer K. 2020. The scientific value of UNESCO biosphere reserves. S Afr J Sci. 116(1/2) https://sajs.co.za/article/view/7432. 10.17159/sajs.2020/7432

Vromans DC, Adams JB, Riddin T. 2013. The phenology of *Ruppia cirrhosa* (Petagna) Grande and Chara sp. in a small temporarily open/closed estuary, South Africa. Aquat Bot. 110:1–5 https://linkinghub.elsevier.com/retrieve/pii/S0304377013000211. 10.1016/j.aquabot.2013.01.008

Mormile JE, Hill CM. 2017. Living With Urban Baboons: Exploring Attitudes and Their Implications for Local Baboon Conservation and Management in Knysna, South Africa. Hum Dimens Wildl. 22(2):99–109 https://www.tandfonline.com/doi/full/10.1080/10871209.2016.1255919. 10.1080/10871209.2016.1255919

Slater K, Barrett A, Brown LR. 2018. Home range utilization by chacma baboon (Papio ursinus) troops on Suikerbosrand Nature Reserve, South Africa Yue B-S, editor. PLoS One. 13(3):e0194717 https://dx.plos.org/10.1371/journal.pone.0194717. 10.1371/journal.pone.0194717

Rosas-Chavoya M et al. 2022. QGIS a constantly growing free and open-source geospatial software contributing to scientific development. Cuad Investig Geográfica. 48(1):197–213 https://publicaciones.unirioja.es/ojs/index.php/cig/article/view/5143. 10.18172/cig.5143

Wallace GE, Hill CM. 2012. Crop Damage by Primates: Quantifying the Key Parameters of Crop-Raiding Events Brockman DK, editor. PLoS One. 7(10):e46636 https://dx.plos.org/10.1371/journal.pone.0046636. 10.1371/journal.pone.0046636

Gilby IC, Pokempner AA, Wrangham RW. 2010. A Direct Comparison of Scan and Focal Sampling Methods for Measuring Wild Chimpanzee Feeding Behaviour. Folia Primatol. 81(5):254–264 https://brill.com/view/journals/ijfp/81/5/article-p254_2.xml. 10.1159/000322354

Amato KR, Van Belle S, Wilkinson B. 2013. A Comparison of Scan and Focal Sampling for the Description of Wild Primate Activity, Diet and Intragroup Spatial Relationships. Folia Primatol. 84(2):87–101 https://brill.com/view/journals/ijfp/84/2/article-p87_4.xml. 10.1159/000348305

Sueur C, Petit O. 2008. Organization of Group Members at Departure Is Driven by Social Structure in Macaca. Int J Primatol. 29(4):1085–1098 http://link.springer.com/10.1007/s10764-008-9262-9. 10.1007/s10764-008-9262-9

Friard O, Gamba M. 2016. BORIS: a free, versatile open-source event-logging software for video/audio coding and live observations Fitzjohn R, editor. Methods Ecol Evol. 7(11):1325–1330 https://besjournals.onlinelibrary.wiley.com/doi/10.1111/2041-210X.12584. 10.1111/2041-210X.12584

Team RC. 2024. R: A language and environment for statistical computing. R Foundation for Statistical Computing.

Fox J, Weisberg S, Price B. 2001. car: Companion to Applied Regression. CRAN Contrib Packag. [published online ahead of print] https://cran.r-project.org/package=car. 10.32614/CRAN.package.car

Ogle DH, Doll JC, Wheeler AP, Dinno A. 2015. FSA: Simple Fisheries Stock Assessment Methods. CRAN Contrib Packag. [published online ahead of print] https://cran.r-project.org/package=FSA. 10.32614/CRAN.package.FSA

Gilbert MF. 2017. Human-olive baboon (Papio anubis) conflicts in farms around Mgori forest reserve, Sindiga, Tanzania. Kenyatta University.

Altmann J, Muruthi P. 1988. Differences in daily life between semiprovisioned and wild-feeding baboons. Am J Primatol. 15(3):213–221 https://onlinelibrary.wiley.com/doi/10.1002/ajp.1350150304. 10.1002/ajp.1350150304

Barrett LP, Stanton LA, Benson-Amram S. 2019. The cognition of ‘nuisance’ species. Anim Behav. 147:167–177 https://linkinghub.elsevier.com/retrieve/pii/S0003347218301477. 10.1016/j.anbehav.2018.05.005

Goodall J. 1986. Social rejection, exclusion, and shunning among the Gombe chimpanzees. Ethol Sociobiol. 7(3–4):227–236 https://linkinghub.elsevier.com/retrieve/pii/0162309586900506. 10.1016/0162-3095(86)90050-6

Balcomb SR, Chapman CA, Wrangham RW. 2000. Relationship between chimpanzee (Pan troglodytes) density and large, fleshy-fruit tree density: Conservation implications. Am J Primatol. 51(3):197–203 https://onlinelibrary.wiley.com/doi/10.1002/1098-2345(200007)51:3%3C197::AID-AJP4%3E3.0.CO;2-C. 10.1002/1098-2345(200007)51:3<197::AID-AJP4>3.0.CO;2-C

Oates AGD & JF. 1994. Colobine Monkeys: Their Ecology, Behaviour and Evolution. 1st (1994). Cambridge University Press.

Colmenares F. 1991. Greeting, aggression, and coalitions between male baboons: Demographic correlates. Primates. 32(4):453–463 http://link.springer.com/10.1007/BF02381936. 10.1007/BF02381936

Patzelt A et al. 2014. Male tolerance and male–male bonds in a multilevel primate society. Proc Natl Acad Sci. 111(41):14740–14745 https://pnas.org/doi/full/10.1073/pnas.1405811111. 10.1073/pnas.1405811111

Cerda-Molina AL et al. 2023. Testing the Challenge Hypothesis in Stumptail Macaque Males: The Role of Testosterone and Glucocorticoid Metabolites in Aggressive and Mating Behavior. Biology (Basel). 12(6):813 https://www.mdpi.com/2079-7737/12/6/813. 10.3390/biology12060813

McLennan MR, Hill CM. 2012. Troublesome neighbours: Changing attitudes towards chimpanzees (Pan troglodytes) in a human-dominated landscape in Uganda. J Nat Conserv. 20(4):219–227 https://linkinghub.elsevier.com/retrieve/pii/S1617138112000349. 10.1016/j.jnc.2012.03.002

John G. Fleagle. 2015. Primate Adaptation and Evolution . Third Edition. By John G. Fleagle. Academic Press. Amsterdam (The Netherlands) and Boston (Massachusetts): Elsevier. x + 441 p.; ill.; index. ISBN: 978-0-12-378632-6. 2013. Q Rev Biol. 90(3):335–336 https://www.journals.uchicago.edu/doi/10.1086/682622. 10.1086/682622

Kaburu SSK et al. 2019. Interactions with humans impose time constraints on urban-dwelling rhesus macaques (*Macaca mulatta*). Behaviour. 156(12):1255–1282 https://brill.com/view/journals/beh/156/12/article-p1255_4.xml. 10.1163/1568539X-00003565

Jayapali U, Perera P, Cresswell J, Dayawansa N. 2023. Does Anthropogenic Influence on Habitats Alter the Activity Budget and Home Range Size of Toque Macaques (*Macaca sinica*)? Insight into the Human-Macaque Conflict. Trees, For People. 13:100412 https://linkinghub.elsevier.com/retrieve/pii/S2666719323000444. 10.1016/j.tfp.2023.100412

Kappeler PM, Barrett L, Blumstein DT, Clutton-Brock TH. 2013. Constraints and flexibility in mammalian social behaviour: introduction and synthesis. Philos Trans R Soc B Biol Sci. 368(1618):20120337 https://royalsocietypublishing.org/doi/10.1098/rstb.2012.0337. 10.1098/rstb.2012.0337

Schradin C. 2013. Intraspecific variation in social organization by genetic variation, developmental plasticity, social flexibility or entirely extrinsic factors. Philos Trans R Soc B Biol Sci. 368(1618):20120346 https://royalsocietypublishing.org/doi/10.1098/rstb.2012.0346. 10.1098/rstb.2012.0346

Hammond P, Lewis-Bevan L, Biro D, Carvalho S. 2022. Risk perception and terrestriality in primates: A quasi-experiment through habituation of chacma baboons (Papio ursinus) in Gorongosa National Park, Mozambique. Am J Biol Anthropol. 179(1):48–59 https://onlinelibrary.wiley.com/doi/10.1002/ajpa.24567. 10.1002/ajpa.24567

Bersacola E, Hill CM, Hockings KJ. 2021. Chimpanzees balance resources and risk in an anthropogenic landscape of fear. Sci Rep. 11(1):4569 https://www.nature.com/articles/s41598-021-83852-3. 10.1038/s41598-021-83852-3

Bolt LM et al. 2025. Edge effects and social behavior in three platyrrhines. Am J Primatol. 87(1) https://onlinelibrary.wiley.com/doi/10.1002/ajp.23610. 10.1002/ajp.23610

Strum SC. 2010. The Development of Primate Raiding: Implications for Management and Conservation. Int J Primatol. 31(1):133–156 http://link.springer.com/10.1007/s10764-009-9387-5. 10.1007/s10764-009-9387-5

