## Supplementary material for "Influence of anthropogenic environments on the activity patterns and vigilance of Cape chacma baboons (*Papio ursinus ursinus*) in the Garden Route, South Africa": Supplemenary Tables

#### ***Supplementary Table 1: Ethogram of the chacma baboons' behaviours used for the Boris analysis***

| **Activity** | **Description** | **Subcategory** |
| --- | --- | --- |
| **Locomotion** | Non-social directional movement | Walking; Running; Climbing; Flight: human, baboon avoidance, other animals, cars |
| **Feeding** | Foraging and ingesting food (chewing, ingesting, handling food or water with hands, feet, or mouth) | Human-derived food; Natural food; Water |
| **Resting** | In a stationary state, the individual is inactive, lying down or asleep | Alone; In-group; Self-grooming |
| **Affiliative** | Grooming, social play, lip-smacking, sweet-grunt, clasping, parents with baby | Grooming; Social-play; Lipsmaking; Sweet-grunt; Clasping; Mother with baby |
| **Agonistic** | Aggression (pushing, hitting, biting, grabbing); Threat (silent or vocalized facial displays, rushing forward, tapping the ground, jumping and chasing, barking, growling, roaring) | Visual threat; Sound threat; Attack |
| **Sexual** | Anogenital presentation; Mount | Anogenital presentation; Mount |
| **Vigilance** | Eyes open and head held high: the focus should not be on another individual in the group or a food source | Vig0: slow glance with a passive position and routine scan; Vig1: repeated glances with a passive position and intense scanning; Vig2: individual on these four legs with an active position and routine scan; Vig3: individual on these two hind legs achieving a glance from both sides with an active and intense stance |

**Supplementary Table 2: Results of binomial generalized linear models (GLMs) testing environmental predictors of chacma baboon behaviours.** Each model evaluated the probability of observing one of five focal behaviours—Affiliation, Feeding, Locomotion, Resting, and Vigilance—in a one-vs-all framework (1 = behaviour present, 0 = absent). Predictors included Habitat (open vs. non-open), Weather (sunny vs. other), and mean number of cars per scan (continuous). Troup identity was included as a categorical covariate to control for inter-group variation but is not displayed. The table reports model coefficients (Estimate β), standard errors (SE), Wald z-values, and two-tailed p-values for each predictor–behaviour combination. Positive coefficients indicate an increased probability of the behaviour occurring under the specified condition; negative coefficients indicate a decreased probability. Statistically significant results (p < 0.05) are highlighted in bold in the table.

| **Behaviour** | **Predictor** | **Estimate (β)** | **SE** | **Wald z** | **P-value** |
| --- | --- | --- | --- | --- | --- |
| Affiliation | Habitat_Open | -0.37385 | 0.15194 | -2.46049 | **0.013875** |
| Affiliation | Weather_Sunny | -0 .38389 | 0.099844 | -3.8449 | **0.000121** |
| Affiliation | mean_cars | -0.00221 | 0.039571 | -0.05594 | 0.955389 |
| Feeding | Habitat_Open | 0.595251 | 0.154692 | 3.847983 | **0.000119** |
| Feeding | Weather_Sunny | 0.262185 | 0.089804 | 2.919532 | **0.003506** |
| Feeding | mean_cars | -0.1649 | 0.032577 | -5.06202 | **4.15E-07** |
| Locomotion | Habitat_Open | -0.3615 | 0.151181 | -2.39117 | **0.016795** |
| Locomotion | Weather_Sunny | 0.094506 | 0.106427 | 0.887988 | 0.374547 |
| Locomotion | mean_cars | -0.19955 | 0.040635 | -4.91089 | **9.07E-07** |
| Resting | Habitat_Open | -0.23722 | 0.162952 | -1.45576 | 0.145458 |
| Resting | Weather_Sunny | 0.404574 | 0.120148 | 3.367305 | **0.000759** |
| Resting | mean_cars | -0.10211 | 0.042062 | -2.4276 | **0.015199** |
| Vigilance | Habitat_Open | 0.144679 | 0.116324 | 1.243763 | 0.213587 |
| Vigilance | Weather_Sunny | -0.17573 | 0.0719 | -2.4441 | **0.014522** |
| Vigilance | mean_cars | 0.235614 | 0.026898 | 8.759597 | **1.96E-18** |

**Supplementary Table 3. Results of binomial generalized linear models (GLMs) testing environmental predictors of chacma baboon behaviours in the female-only dataset.** Each model evaluated the probability of observing one of five focal behaviours, Affiliation, Feeding, Locomotion, Resting, and Vigilance, using a one-vs-all framework (1 = behaviour present, 0 = absent). Predictors included Habitat (open vs. non-open), Weather (sunny vs. other), and mean number of cars per scan (continuous). Troup identity was included as a categorical covariate to control for inter-group variation but is not displayed. The table reports model coefficients (Estimate β), standard errors (SE), Wald z-values, and two-tailed p-values for each predictor–behaviour combination. Positive coefficients indicate a higher probability of the behaviour under the specified condition; negative coefficients indicate a lower probability. Statistically significant results (p < 0.05) are shown in bold.

| **Behaviour** | **Predictor** | **Estimate (β)** | **SE** | **Wald z** | **P-value** |
| --- | --- | --- | --- | --- | --- |
| Affiliation | Habitat_Open | -0.07194 | 0.258072 | -0.27877 | 0.780423 |
| Affiliation | Weather_Sunny | -0.57516 | 0.164298 | -3.50069 | **0.000464** |
| Affiliation | mean_cars | 0.105198 | 0.040475 | 2.599099 | **0.009347** |
| Feeding | Habitat_Open | 1.174009 | 0.258585 | 4.54013 | **5.62E-06** |
| Feeding | Weather_Sunny | 0.178444 | 0.149932 | 1.190168 | 0.233981 |
| Feeding | mean_cars | -0.06535 | 0.036165 | -1.80696 | 0.070769 |
| Locomotion | Habitat_Open | -0.3742 | 0.221018 | -1.69309 | 0.090439 |
| Locomotion | Weather_Sunny | -0.0081 | 0.165571 | -0.04894 | 0.960971 |
| Locomotion | mean_cars | -0.07115 | 0.042032 | -1.69284 | 0.090485 |
| Resting | Habitat_Open | -0.60068 | 0.219806 | -2.73277 | **0.00628** |
| Resting | Weather_Sunny | 0.501421 | 0.188915 | 2.654215 | **0.007949** |
| Resting | mean_cars | 0.033852 | 0.042134 | 0.80344 | 0.421721 |
| Vigilance | Habitat_Open | -1.54913 | 1.292783 | -1.19829 | 0.230804 |
| Vigilance | Weather_Sunny | -1.22466 | 1.187002 | -1.03173 | 0.3022 |
| Vigilance | mean_cars | -0.59095 | 0.843215 | -0.70082 | 0.483413 |
